# Lipid A phosphoethanolamine transferase-mediated site-selective modifications show association with colistin resistance phenotypes and fitness

**DOI:** 10.1101/2024.08.20.608901

**Authors:** A. Schumann, A. Gaballa, H. Yang, D. Vu, RK. Ernst, M. Wiedmann

## Abstract

Genes encoding lipid A modifying phosphoethanolamine transferases (PETs) are genetically diverse and can confer resistance to colistin and antimicrobial peptides. To better understand the functional diversity of PETs, we characterized three canonical mobile colistin resistance (*mcr*) alleles (*mcr-1*, *-3*, *-9*), one intrinsic *pet* (*eptA*), and two *mcr*-like genes (*petB*, *petC*). Using an isogenic expression system, we show that *mcr-1* and *mcr-3* are phenotypically similar by conferring colistin resistance with lower fitness costs. *mcr-9*, which is phylogenetically closely related to *mcr-3*, and *eptA* only provide fitness advantages in the presence of sub-inhibitory concentrations of colistin and significantly reduce fitness in media only. PET-B and PET-C were phenotypically distinct from bonafide PETs; neither conferred colistin resistance or caused considerable fitness cost in *Escherichia coli*. Strikingly, we found for the first time that different PETs selectively modify different phosphates of lipid A - MCR-1, MCR-3, and PET-C selectively modify the 4’-phosphate, while MCR-9 and EptA modify the 1-phosphate. 4’-phosphate modifications facilitated by MCR-1 and -3 are associated with high levels of colistin resistance and low toxicity. Our results suggest that PETs have a wide phenotypic diversity and that high level colistin resistance is associated with specific lipid A modification patterns that has been largely unexplored thus far.

**IMPORTANCE:** Rising levels of resistance to increasing numbers of antimicrobials has led to the revival of last resort antibiotic colistin. Unfortunately, resistance to colistin is also spreading in the form of *mcr* genes, making it essential to (i) improve identification of resistant bacteria to allow clinicians to prescribe effective drug regimens and (ii) develop new combination therapies effective at targeting resistant bacteria. Our results demonstrate that PETs, including MCR variants, are site-selective in *E. coli*, with site-selectivity correlating with the level of resistance and fitness costs conferred by certain PETs. Site-selectivity associated with a given PET may not only help predict colistin resistance phenotypes, but may also provide an avenue to (i) improved drug regimens and (ii) development of new combination therapies to better combat colistin resistant bacteria.

## INTRODUCTION

Gram-negative bacteria are becoming more resistant to available antimicrobials. Hence, last-resort antimicrobials including colistin (polymyxin E) are increasingly used as therapeutics (1, 2). Unfortunately, colistin resistance (col^R^) has been spreading through mobile colistin resistance (*mcr*) genes (3). MCR variants encoded by these genes are part of enzymes specific to Gram-negative bacteria, classified as lipid A phosphoethanolamine transferases (PETs); that catalyze the transfer of phosphoethanolamine (pEtN) to one of the terminal phosphate moieties of lipid A (4, 5). This modification reduces the overall negative charge of the outer membrane, decreasing colistin’s ability to interact with the negatively charged phosphates of lipid A (6). To date, ten different MCR families have been discovered that share varying degrees of amino acid sequence similarity with themselves and the intrinsic PET, EptA (3, 4). While MCR variants are well known for providing col^R^, recent studies have reported that MCR expression can lead to fitness costs (7–9), potential cross-resistance to other antimicrobial peptides (AMPs) (10–12), or impact bacterial-host interactions (10, 13). However, the functional diversity of MCR and PET variants is scarcely understood.

To quantitatively probe the phenotypic diversity of MCR variants, as well as PETs that have been linked to the stress response to AMPs (14), we optimized a standardized and monitorable expression system to physiologically and biochemically characterize a range of canonical (i.e., EptA, MCR-1, -3, -9) and novel PETs (i.e., PET-B, -C). We found that PET families represent diverse phenotypes and that PETs show stereoselective pEtN modifications of lipid A in *Escherichia coli*, which may be associated with col^R^ phenotypes and may provide avenues for the development of new antibiotic adjuvants and antimicrobial drugs.

## RESULTS

### Cell viability defects and colistin MICs of canonical PETs differ

To better characterize the phenotypic diversity of PETs, we created a standardized expression system that includes the colistin susceptible strain *E. coli* Top10 (15) and vector pBAD24 with an L-arabinose-inducible promoter (4). Four canonical *pet* genes (i.e., *mcr-1*, *mcr-3*, *mcr-9*, and *eptA*) were cloned into this expression system, and protein expression was confirmed for uninduced and induced strains using western blots (Fig. 1A, Fig. S1). While MCR-1 and MCR-3 protein levels were significantly higher when induced with either 0.05 or 0.2% of L-arabinose (p<0.01), there was no significant difference in EptA and MCR-9 protein levels between induced and uninduced strains (Fig. 1A). Interestingly, MCR-9 protein levels were significantly lower than MCR-1 levels for samples induced with 0.05 or 0.2% L-arabinose (p<0.01) and MCR-3 levels for samples induced with 0.2% L-arabinose (p<0.05) (Fig. 1a).

**FIG 1.**
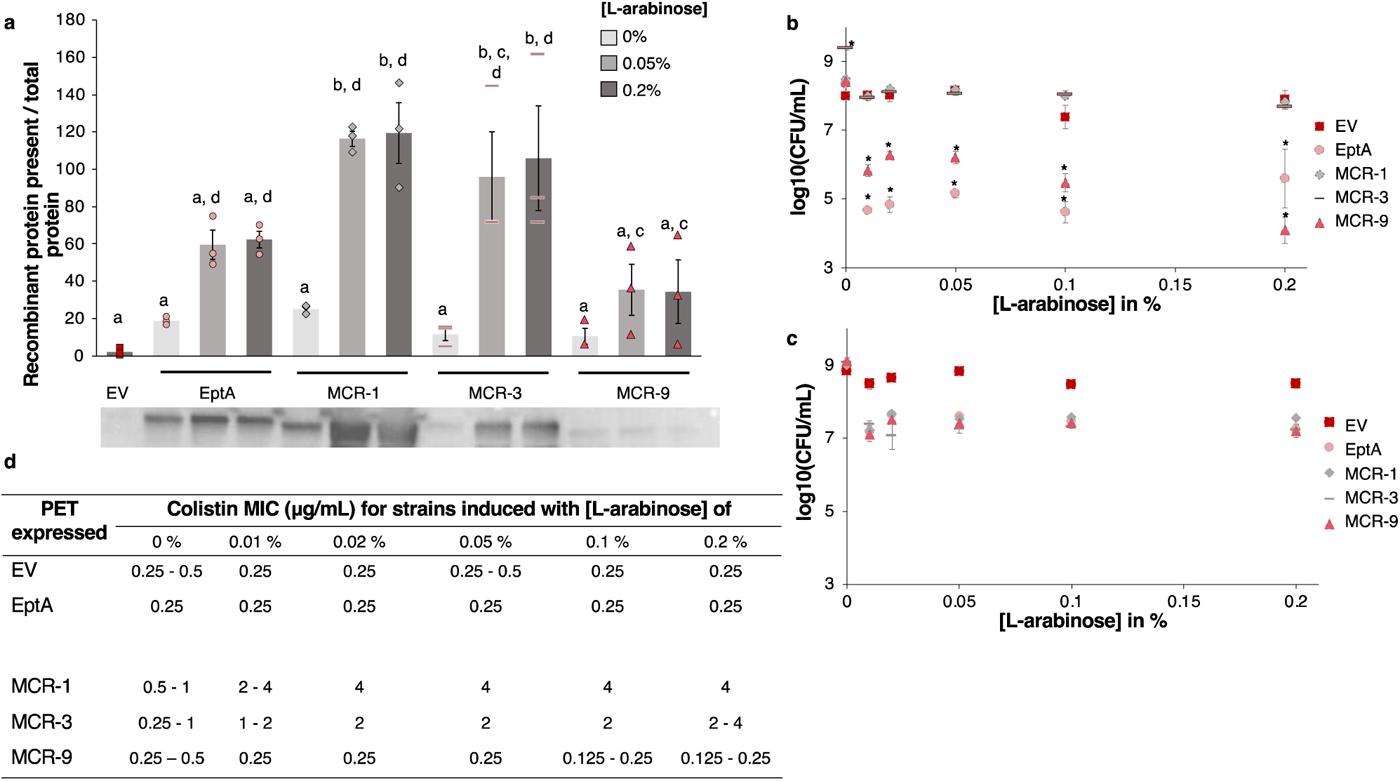
Characteristics of *E. coli* Top10 expressing canonical PET variants induced with different L-arabinose concentrations. For all experiments, strains were grown in MHII media for 2 hrs at 37°C, 200 rpm prior to induction. (**A**) Western blot of cells collected after 12 hrs of induction with 0.05 or 0.2% of L-arabinose; anti-FLAG antibodies were used to detect heterologously expressed FLAG-tagged PET proteins, data represent means, n = 3 ± SEM, samples with the same letter are not significantly different, using a one-way ANOVA and post-hoc Tukey test. (**B, C**) Cell viability assessment after 4 (**B**) and 12 hrs (**c**) of incubation with different L-arabinose concentrations, data represent means, n = 3 ± SEM, * indicates significant differences from the EV control at the same concentration (α=0.05) using emmeans after a two-way ANOVA. (**D**) Colistin MICs of strains added to plates after 1 hr of induction with different L-arabinose concentrations; MICs were determined from biological triplicates; two concentrations mean strains had two different MICs.

To identify an optimal L-arabinose concentration to induce PET expression, we measured cell viability and colistin Minimum Inhibitory Concentrations (MICs) for strains expressing canonical PETs, in comparison to an empty vector control (“EV”) at different inducer concentrations (Fig. 1, Fig. S2). After induction for 4 hours, the cell viability of strains expressing MCR-1 or -3 was not significantly different from the EV across all inducer concentrations (Fig. 1B, Fig. S2). In contrast, strains expressing EptA or MCR-9 showed significant reductions in cell viability (relative to the EV) across all inducer concentrations (p<0.05); for example, MCR-9 and EptA expressing strains showed mean cell viability reductions of 1.9 and 3.0 log, respectively, when induced with 0.05% L-arabinose (Fig. 1B).

Inducer concentrations also impacted colistin MICs. For all strains, 0.02 or 0.05% L-arabinose yielded maximum MIC levels, which were (i) 0.25 μg/mL for the EV, EptA, and MCR-9, (ii) 2 μg/mL for MCR-3, and (iii) 4 μg/mL for MCR-1 expressing strains (.1D). Hence, we selected 0.05% L-arabinose as our standardized expression condition for further experiments because this concentration resulted in (i) maximum col^R^, (ii) minimum cell viability defects, and (iii) reliable protein expression across all our PETs (Fig. 1).

### Novel *mcr*-like PETs are phenotypically distinct from canonical PETs

We previously predicted novel *mcr*-like genes that may constitute new MCR families (4). To further characterize the diversity of PETs, we characterized two of those predicted putative, novel *mcr*-like PETs (i.e., PET-B, -C) (Fig. 2A). Western blot analysis confirmed that both proteins were expressed (Fig. S3). Colistin MICs of PET-B and -C in *E. coli* were similar when uninduced and induced (0.05% L-arabinose) showing MICs of 0.25 – 0.5 μg/mL (Fig. 2B). We however found evidence the expression of PET-B and - C conferred some cytotoxicity; compared to uninduced controls; strains expressing PET-B showed mean log reductions in cell viability of 0.8 and 0.7 after 12- and 22-hours, while PET-C showed 0.8 and 1.2 log reductions after 4- and 12-hours of induction (p<0.05) (Fig. 2C).

**FIG 2.**
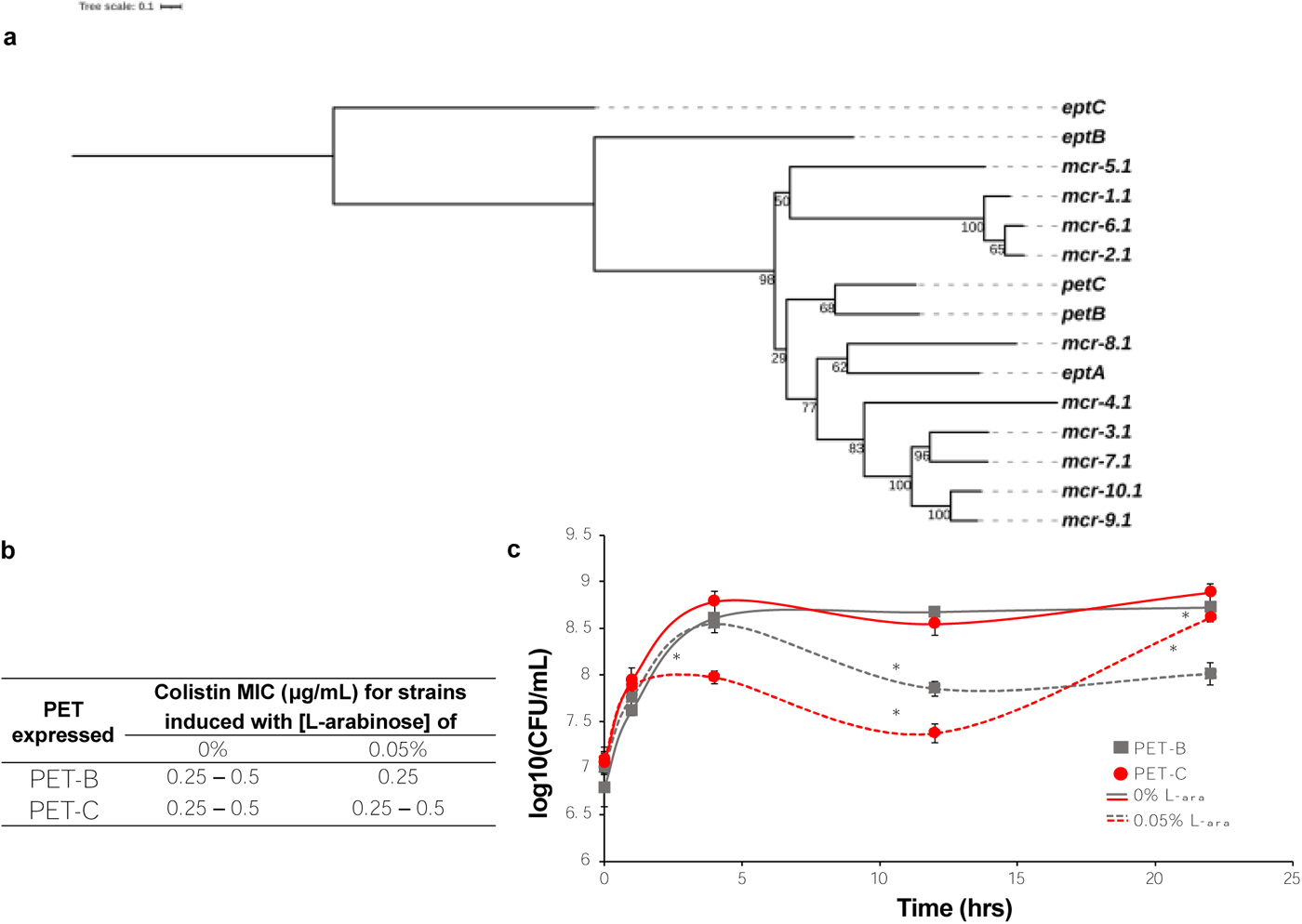
Novel, *mcr-*like PETs in *E coli* Top10. (**A**) Phylogenetic tree shows the relatedness of novel, *mcr*-like PETs to chromosomally encoded PETs (i.e., *eptA*, *eptB*, and *eptC*) and *mcr* variants. (**B**) Colistin MICs were obtained after growth in MHII broth for 2 hrs, followed by induction with 0 or 0.05% of L-arabinose for 1 hr. MICs were determined from biological triplicates; two concentrations mean strains had two different MICs. (**C**) Cell viability was determined at several time points after induction with 0 or 0.05% of L-arabinose, data represents means, n = 3 ± SEM, * indicates significant differences between the uninduced, isogenic control (α=0.05) based on emmeans after a two-way ANOVA.

### PET-expressing strains differ in their response to various cell envelope stressors

As MCR-1 has been shown to provide cross-resistance to AMPs (10, 12, 16), we characterized the ability of different PET-expressing strains to confer resistance to different AMPs and cell envelope stressors. Strains expressing EptA or MCR-9 showed considerably lower EDTA (a metal chelator) MICs (4 and 2 μg/mL) than strains expressing any other PET or the EV (MIC of 16 μg/mL) (Table 1). In zone of inhibition (ZOI) assays, colistin, and cecropin A, a positively charged AMP (17), were the only compounds that yielded visible ZOIs (Table 2). While the EV had the largest ZOI for colistin and cecropin A, ZOIs did not substantially differ between PET-expressing strains (e.g., cecropin A ZOIs ranged from 11 to 12 mm).

**TABLE 1.**
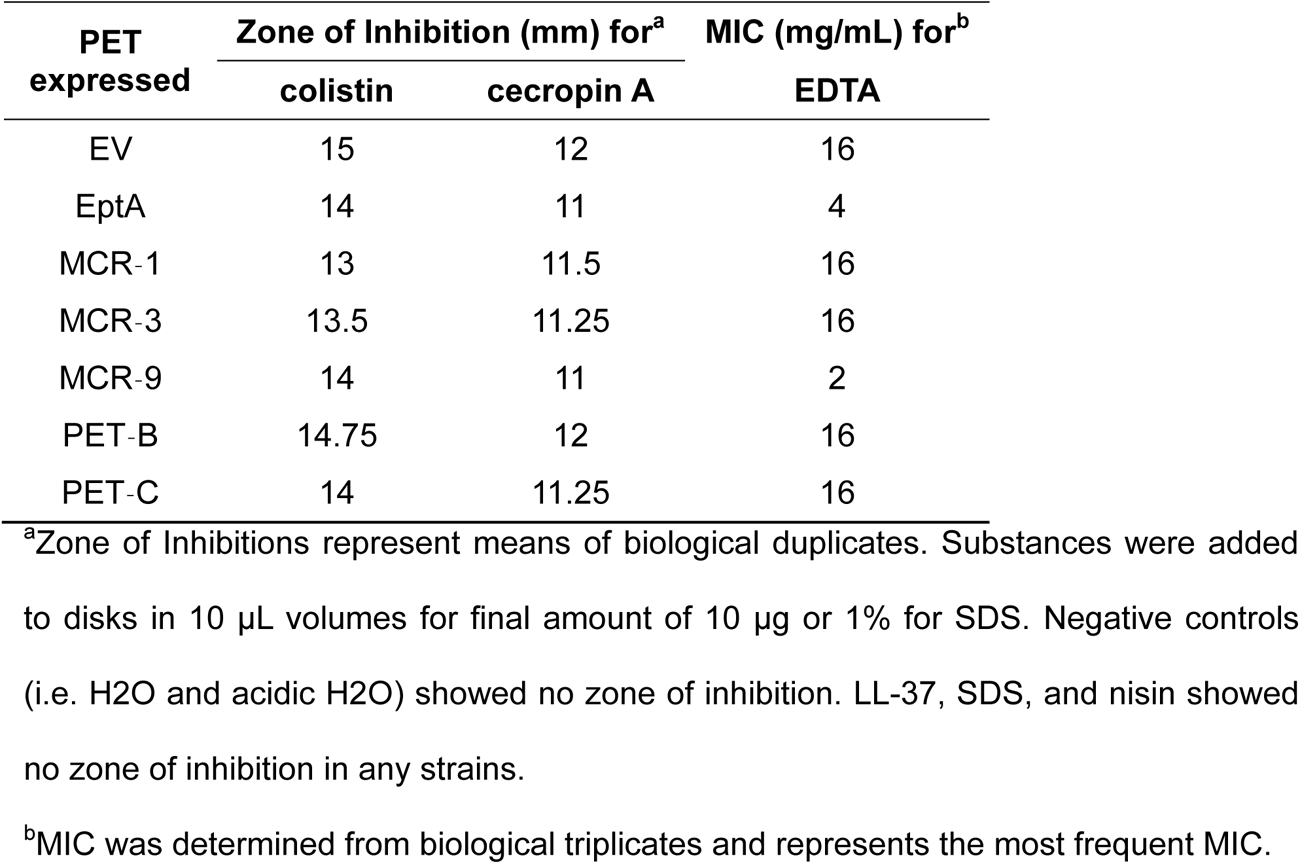
Zone of Inhibitions and MIC values of *E. coli* Top10 strains expressing PET variants, induced with 0.05% L-arabinose to antimicrobial peptides or cell envelope stressors.

| PET<br>expressed | Zone of Inhibition (mm) for<sup>a</sup> |  | MIC (mg/mL) for<sup>b</sup> |
| --- | --- | --- | --- |
|  | colistin | cecropin A | EDTA |
| EV | 15 | 12 | 16 |
| EptA | 14 | 11 | 4 |
| MCR-1 | 13 | 11.5 | 16 |
| MCR-3 | 13.5 | 11.25 | 16 |
| MCR-9 | 14 | 11 | 2 |
| PET-B | 14.75 | 12 | 16 |
| PET-C | 14 | 11.25 | 16 |

<sup>a</sup>Zone of Inhibitions represent means of biological duplicates. Substances were added to disks in 10 µL volumes for final amount of 10 µg or 1% for SDS. Negative controls (i.e. H<sub>2</sub>O and acidic H<sub>2</sub>O) showed no zone of inhibition. LL-37, SDS, and nisin showed no zone of inhibition in any strains.
<sup>b</sup>MIC was determined from biological triplicates and represents the most frequent MIC.

### PETs confer context-dependent fitness advantages and disadvantages

We further characterized our PET variants by performing (i) one-hour killing assays and (ii) competition assays. Compared to static MIC values, both assays allow for better assessments of the dynamic nature of bacterial interactions with antibiotics (18).

In the killing assays, higher colistin concentrations corresponded to higher log reductions for the EV and all PET*-*expressing strains except MCR-1. For all strains (except MCR-1), the relative log reduction was < 1 log after exposure to 1 and 2 μg/mL colistin but increased to 1.6 - 6.2 after exposure to 8 μg/mL colistin (Fig. 3A). The strain expressing MCR-1 showed consistent col^R^ (< 0.2 log reduction at all tested colistin concentrations). Differences between PETs were most apparent at 8 μg/mL of colistin: both MCR-1 (0.2 log reduction) and MCR-3 (1.6 log reduction) conferred reduced die-off as compared to the EV (2.9 log reduction). While the EptA strain showed a 2.3 log reduction (similar to EV), MCR-9, PET-B, and PET-C showed substantially increased reductions (6.1, 6.2, and 5.4 log, respectively) relative to the EV (Fig. 3A).

**FIG 3.**
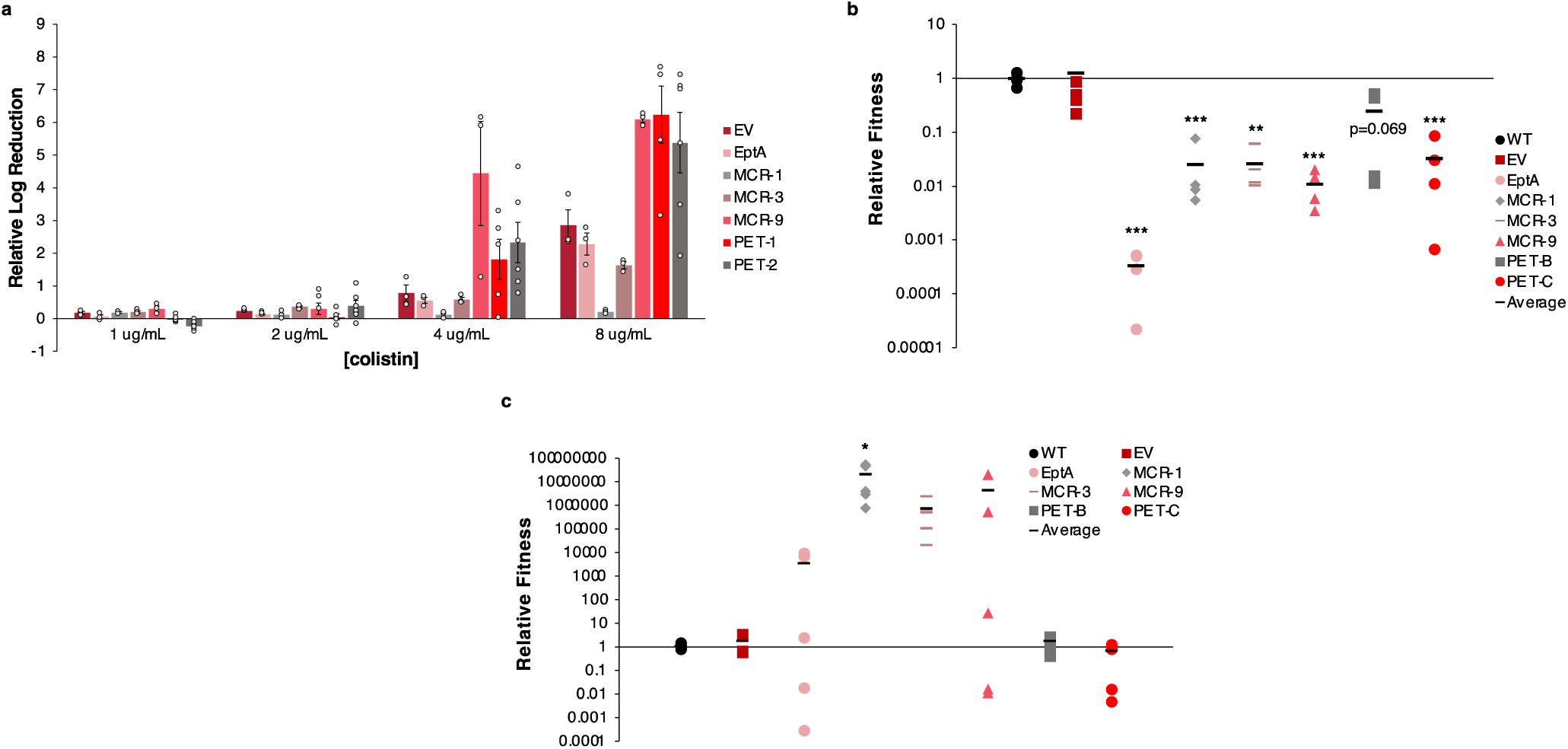
Colistin Resistance and fitness of PET expressing *E. coli* Top10 strains. For all data shown, strains were grown for 2 hours at 37°C and induced with 0.05% L-arabinose for 1 hour prior to experiments. (**A**) Relative log reduction of *pet* expressing strains after exposure to different colistin concentrations in 1 hour killing assays in comparison to strains grown in the absence of colistin, data represent means, n=3 ± SEM, except n=5 (PET-B), n=6 (PET-C). (**B**) Fitness of *pet* expressing strains (*lacZ^-^*) after 12 hours competition with *E. coli* pUC19 (*lacZ^+^*) strains in MHII without colistin. Relative fitness was calculating by adjusting fitness values by the average fitness of the WT strain for each experiment, black bars represent means, n = 4, significance values based on comparison to WT strains using a one-way ANOVA and post hoc Dunnett’s test. (**C**) Fitness of *pet* expressing strains (*lacZ^-^*) after 12 hours competition with *E. coli* pUC19 (*lacZ^+^*) strains in MHII the presence of colistin (concentrations at ¼ of each strain’s colistin MIC). Fitness values are relative to the WT strain, black bars represent means, n = 5, significance values are based on comparisons to the WT strains using a Kruskal-Wallis test and post-hoc Many-to-One Dunn’s test. *, p<0.05; **, p<0.01, ***, p<0.001

Competition assays were used to assess fitness by determining the relative changes in cell numbers of two strains grown in co-cultures (19). Competition assays were performed in the presence and absence of colistin to understand the context-dependent fitness costs of PET expression. Here, fitness is defined as the ratio of surviving PET-expressing cells to surviving competitor cells (*E. coli* Top10 pUC19), normalized by their initial levels. In the absence of colistin, the expression of different PETs conferred varying degrees of fitness costs; strains expressing (i) EptA were recovered at >1000-fold lower levels (p<0.001), (ii) MCR-1, PET-C, MCR-9 (p<0.001), and MCR-3 (p<0.01) were recovered at 10-to 100-fold lower levels, and (iii) PET-B was recovered at <10-fold lower levels (p=0.069) than the competitor strain (Fig. 3B).

*E. coli* expressing canonical PETs were considerably more fit than the competitor strain (Fig. 3C) in the presence of colistin at 1/4^th^ of the strain’s MIC (Fig. 1D, Fig. 2B). Strains expressing EptA were recovered at 100-fold, MCR-1 at 10^7^-fold (p<0.05), MCR-3 and MCR-9 at 10^6^-fold higher levels than the competitor strain, while strains expressing PET-B or -C were recovered to the same extent as the competitor strain (Fig. 3C). Interestingly, the competitive indices of EptA (p<0.01) and MCR-9 (p<0.05) expressing strains varied significantly more when compared to all other strains (except for the comparison between variances for EptA and PET-C expressing strains).

### PET enzymes are stereoselective in their lipid A modification

We confirmed the functionality of our heterologous proteins by assessing pEtN-modification of lipid A 12 hours after induction with 0.05% L-arabinose. The detected ions at *m/z* 1797 and 1920 indicated hexa-acylated lipid A and pEtN-modified hexa-acylated lipid A (Δ123 *m/z*), respectively (Fig. 4A). PEtN modifications were detected for strains expressing EptA, MCR-1, MCR-3, MCR-9, and PET-C, but not for the EV or PET-B expressing strains (Fig. 4A, Table S1).

**FIG 4.**
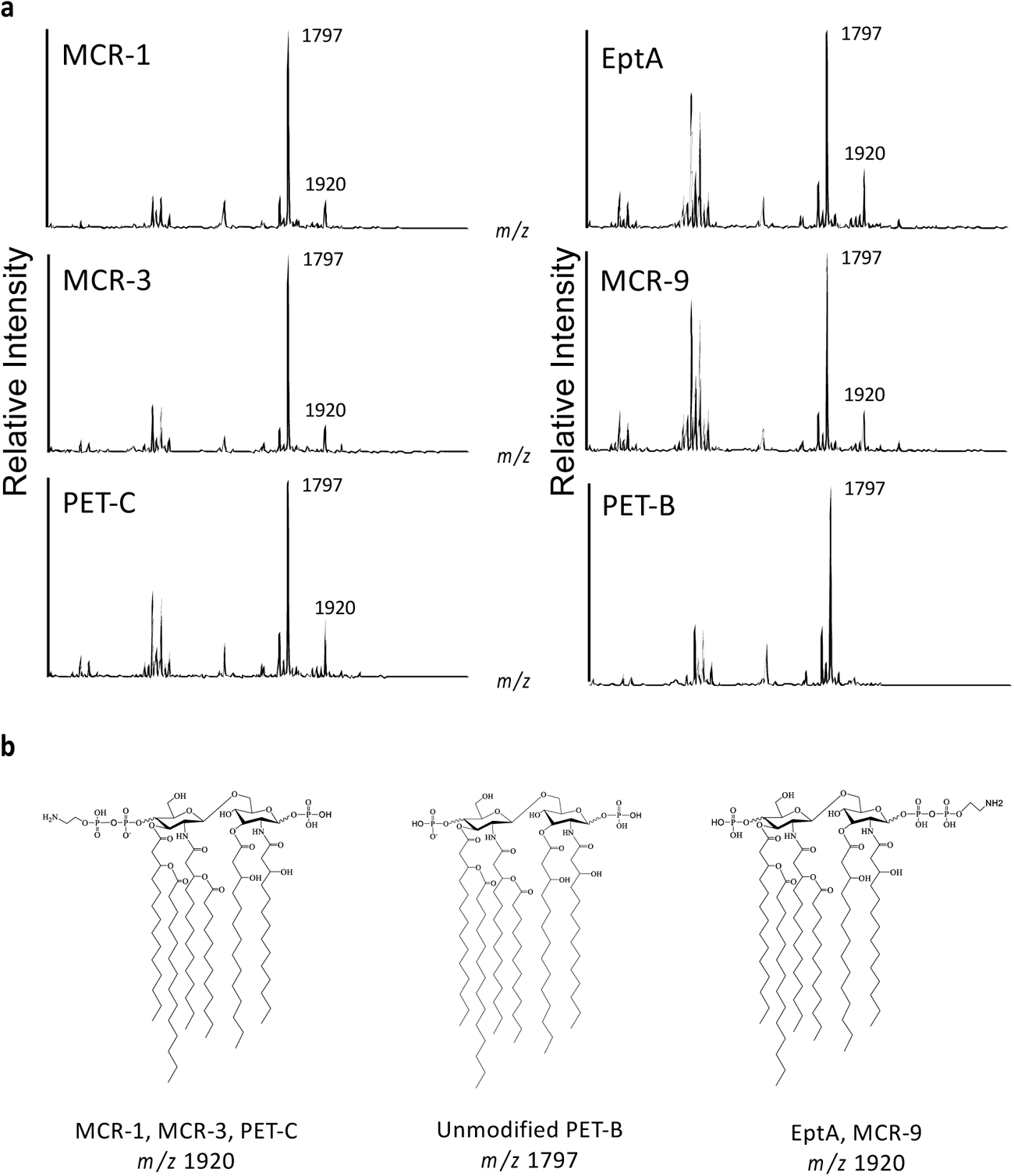
Site-selective modification of lipid A by PETs. FLAT mass spectra and chemical structure of lipid A. (**A**) Six different strains (MCR-1, MCR-3, PET-C, EptA, and MCR-9) show lipid A signals at *m/z* 1797 and 1920, while PET-B only shows a lipid A signal at *m/z* 1797. (**B**) Chemical structure of each lipid A; while MCR-1, MCR-3, and PET-C have 4’-phosphate group modification with pEtN, the others, EptA and MCR-9, have 1-phosphate group modification. PET-B does not show pEtN modification. All structural analyses were conducted by tandem MS (FLAT^n^), as shown in Fig. 5.

To determine the stereospecificity of individual PETs, tandem mass spectra were generated with MS/MS patterns showing two structural isomers from the precursor ion at *m/z* 1919.22, allowing for the determination of the pEtN location (20). MCR-1, MCR-3, and PET-C showed a fragmentation pattern with a base peak at *m/z* 1821.24 and a diagnostic ion of 4’-phosphate modification with pEtN at *m/z* 1267.81 (Fig. 5A). These ions originate from the loss of 1-phosphate and cross-ring cleavage (^0,4^A_2_), respectively (Fig. 5B). In contrast, EptA and MCR-9 showed a fragmentation pattern of the precursor ion at *m/z* 1919.21 with a base peak at *m/z* 1243.83 and a diagnostic ion of 1-phosphate modification with pEtN at *m/z* 833.43 (Fig. 5A). These ions originate from multiple neutral losses of an acyl chain, 4-phosphate, and pEtN (1 + B’2 + 3’α) and glycosidic bond dissociation (Y1), respectively (Fig. 5B). The ratio of intensities of the two diagnostic ions at *m/z* 833.43 (1-phosphate modification) and 1267.81 (4’-phosphate modification) indicated that MCR-1, MCR-3, and PET-C selectively attach the pEtN residue to the 4’-phosphate with 97, 98, and 97% efficacy, respectively. In contrast, EptA and MCR-9 preferentially attach the pEtN residue to the 1-phosphate with 75% efficacy (Fig. S4, Fig. 4B).

**FIG 5.**
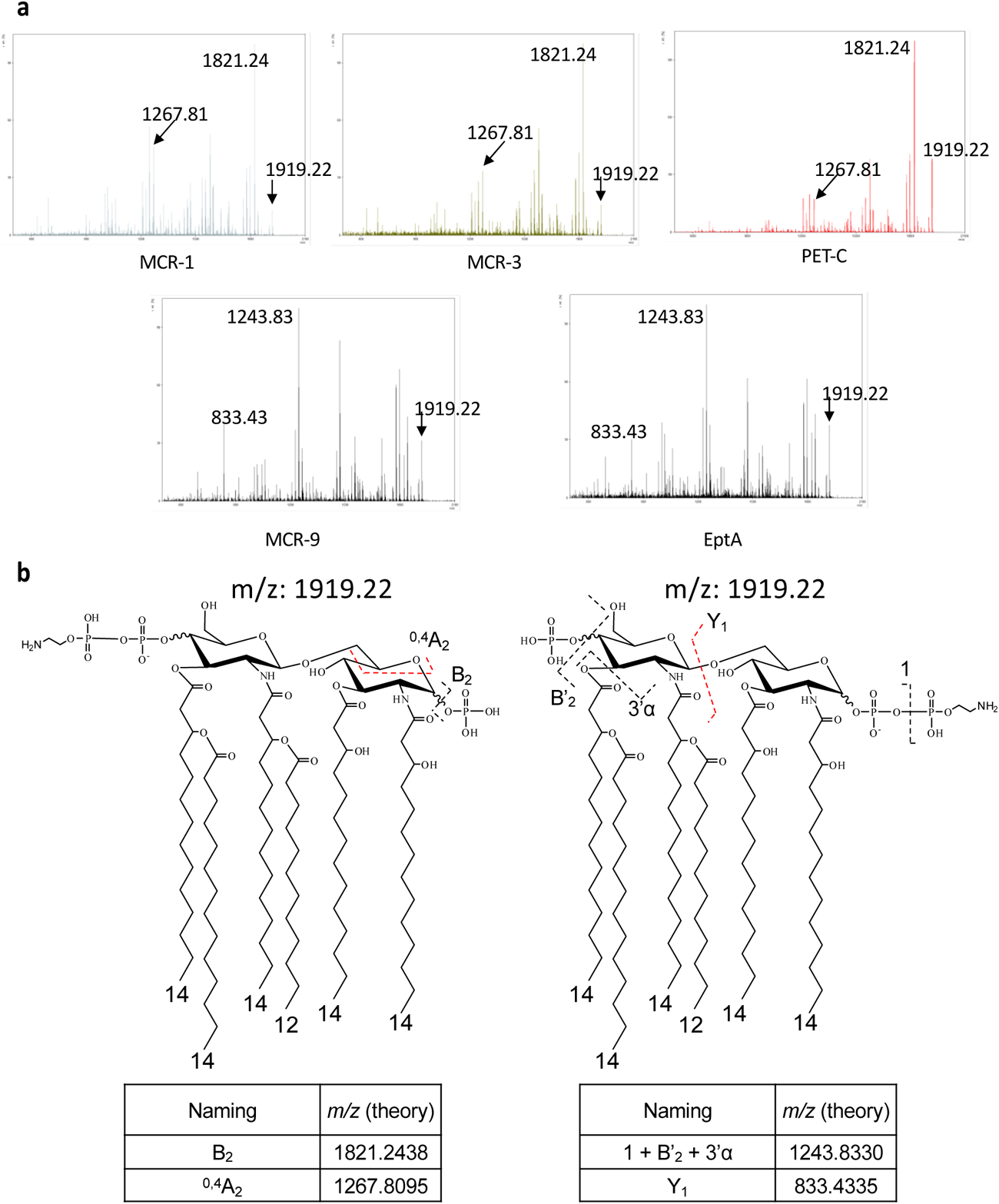
pEtN positions determination by FLATn. FLATn conducted the structural analysis of pEtN-modified lipid A at *m/z* 1919.22. (**A**) Five different PET-expressing strains (MCR-1, MCR-3, PET-C, MCR-9, and EptA) of *E. coli* have been investigated to determine the modification site by pEtN. Three stains (MCR-1, MCR-3, and PET-C) show similar fragmentation patterns each other that have *m/z* 1821.24 as a base peak and diagnostic ion at *m/z* 1267.81, which indicates the modification site by pEtN is 4’-phosphate. On the other hand, two other stains (MCR-9 and EptA) show different fragments compared to MCR-1, MCR-3, and PET-C. The base peak and diagnostic fragment in FLATn mass spectra of MCR-9 and EptA are *m/z* 1243.83 and 833.43, indicating that modification site by pEtN is 1-phosphate. (**B**) Chemical structures of the modified lipid A on the phosphate group with pEtN are shown; the fragments and their theoretical m/z values are listed below the chemical structures.

### PET-B’s extended loop region differs structurally from other PETs

Consistent with previous reports (21), our structural models showed that PETs share a highly conserved overall domain structure with discretely folded N-terminal membrane-anchored and C-terminal soluble periplasmic domains (Fig. 6A, B). The structural similarities dendrogram and the correspondence analysis of the structural similarity matrix showed that PETs formed two distinct groups, including (i) MCR-1, -2, and -6, and (ii) EptA, PET-B, PET-C, and the remaining MCR variants (Fig. S5). The modeled PET-B and -C structures showed similar domain architecture and conserved residues involved in the catalytic and pEtN binding sites (Fig. 6A, B). However, superimposed structural models indicated that structures of a periplasmic loop separating two membrane-anchored alpha-helices, as well as the bridging alpha-helix, and extended loop connecting the membrane-anchored and periplasmic domains, differ substantially between PET-B and -C (Fig. 6C, D). While the bridging helix and extended loop are conserved among PET-C, EptA, MCR-3, and MCR-9 (Fig. 6E), they are markedly different between MCR-1 and other PET-predicted structures (Fig. 6E, Fig. S5).

**FIG 6.**
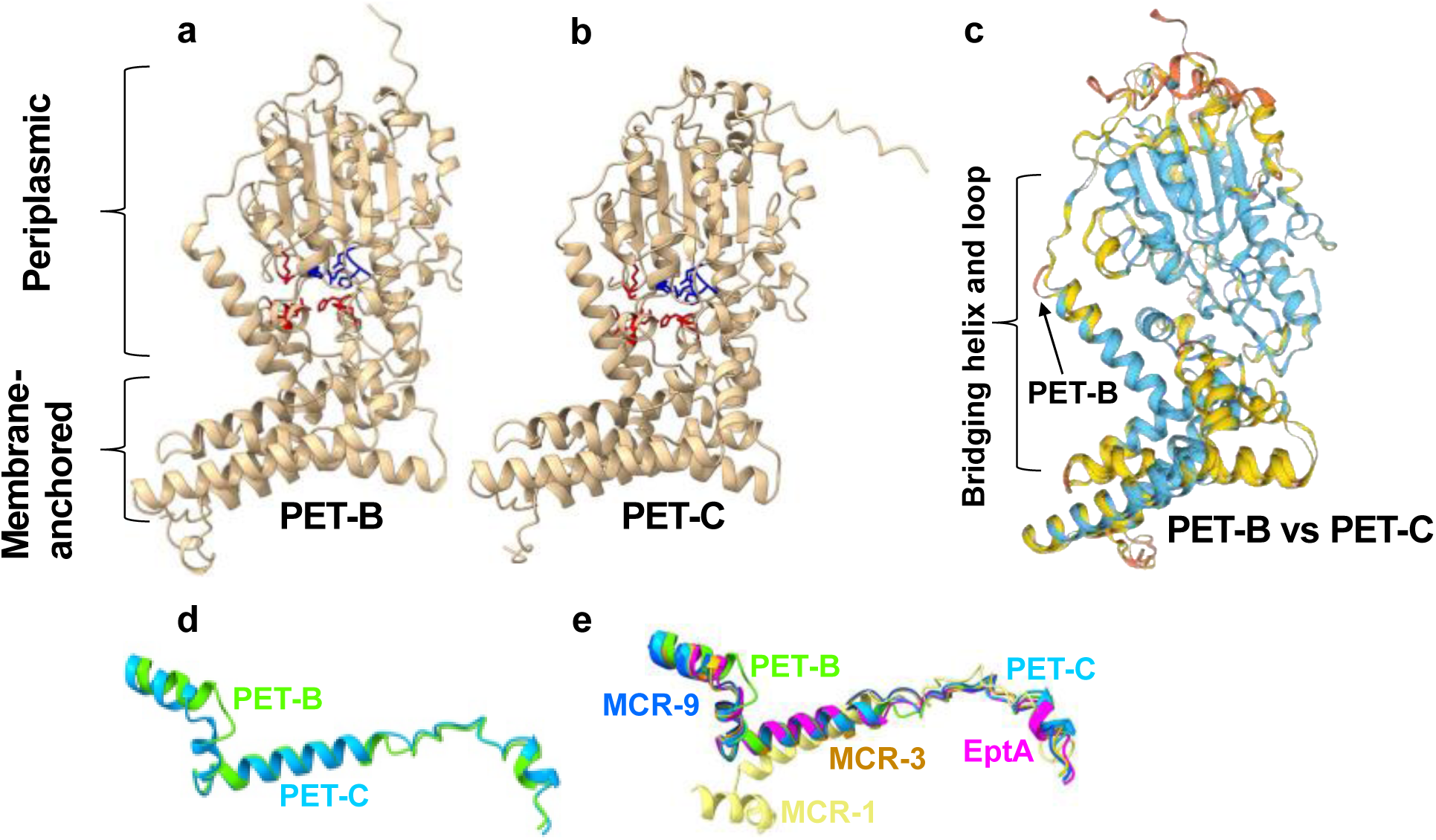
PET-B and PET-C share conserved domain architecture with localized structural variations. (**A, B**) PET-B (**A**) and PET-C (**B**) structural models were predicted using AlphaFold2 based on the *N. meningitidis* phosphoethanolamine transferase EptA structure as a template (NCBI structure accession number 5FGN). Structures were viewed and edited using UCSF ChimeraX. Structural models show the conservation of distinct transmembrane-anchored and soluble periplasmic domain architecture and AA residues involved in forming the catalytic (blue residues) and pEtN binding site (red residues). (**C**) A superimposed image of the PET-B and PET-C structures was constructed using Swiss-Model structural comparison tools, showing regions with low similarity confidence (yellow) and very low similarity confidence (orange). (**D)** Localized structural variability of the bridging helix and extended loop regions of PET-B (light green) and PET-C (light blue). PET-B and PET-C structural comparisons were performed using the Swiss-Model server, and the superimposed structures were downloaded, viewed, and edited using UCSF ChimeraX. (**E**) Localized structural variability of the bridging helix and extended loop regions of PET-B (light green), PET-C (light blue), EptA (magenta), MCR-1 (yellow), MCR-3 (orange) and MCR-9 (blue). Structural comparisons were performed using the Swiss-Model server, and the superimposed structures were downloaded, viewed, and edited using UCSF ChimeraX.

## DISCUSSION

Here, we used a standardized expression system to functionally and biochemically characterize six distinct *pet* genes, including (i) two widely distributed *mcr* variants with a well-supported ability to confer col^R^ (*mcr-1* and *mcr-3*), (ii) *mcr-9*, a *mcr* gene with a disputed ability to confer col^R^; (iii) *eptA*, an intrinsic *pet* involved in the bacterial stress response to AMPs, and (iv) two chromosomally encoded uncharacterized *pet* genes (*petB* and *petC*, referred to as “*mcr*-like PETs”). Our data indicate that known MCR variants (i.e., MCR-1, -3, and -9) differ in their ability to confer col^R^. While expression of MCR variants incurs a range of fitness costs in the absence of colistin, it appears to provide variable fitness advantages in the presence of subinhibitory levels of colistin. We also found that the two *mcr*-like PETs characterized here included one gene (*petC*) that modifies lipid A and one gene (*petB*) that had no detectable phenotype in *E. coli*; these results may be explained by the fact that these genes have primarily been identified in *Pseudomonas* (4), which may have a different lipid A structure and use different lipid A modification enzymes (22, 23). Most strikingly, we found that MCR-1 and -3, which conferred high levels of col^R^, stereoselectively modify lipid A on the 4’-phosphate, while EptA and MCR-9 modify the 1-phosphate.

While we established an optimized expression system with a fine-tuned induction concentration, previous studies using similar expression strains typically used a higher L-arabinose concentration (i.e., 0.2%), which caused greater cellular defects that might impact col^R^ which was not assessed (7, 8, 24). Using our *E. coli* expression system, we found that MCR-1 and -3 conferred the highest resistance levels (i.e., MICs of 4 and 2 μg/mL, respectively), wild-type strains carrying these genes typically showed similar MICs, e.g., MICs of 4-8 μg/mL for *E. coli* carrying either *mcr-1* or *-3* (25, 26). Killing assay data suggested phenotypic differences between MCR-1 and -3, with MCR-1 providing protection at higher colistin concentrations. We also showed that MCR-1 and - 3 expression conferred the lowest fitness costs, which suggests that these two PETs have evolved to provide effective col^R^ with minimal impact on cell growth. This is consistent with previous observations that fitness costs associated with *mcr-1* expression are rarely detected in natural strains with *mcr*-expressing vectors (7, 27) and the fact that *mcr-1* and *-3* represent the most diverse *mcr* families with 37 and 42 variants, respectively (NCBI Pathogen Detection, Reference Catalog, 06/27/24).

Our study also provides new insights on MCR-9, which has been of interest as it is frequently found in isolates classified as colistin susceptible (28). While several studies did not show increased MICs of strains naturally carrying or expressing *mcr-9* (28, 29), Kananizadeh et al. (30) found that *mcr-9*-positive *Enterobacter cloacae* isolates are resistant to colistin when grown in Brain-Heart Infusion, Tryptic Soy, and Lysogenic broth (MICs of 4 to 128 μg/mL), but not when grown in cation-adjusted Müller-Hinton broth (MHII; MICs of 0.125 – 0.5 μg/mL). While we found similar MICs in MHII, in a competition assay, we found that MCR-9 expression conferred fitness advantages in the presence of subinhibitory levels of colistin. While MCR-9 does not seem to confer resistance detectable in standard MIC assays, it does appear to provide fitness advantages in the presence of colistin.

Importantly, our data showed that overexpression of the four canonical PETs resulted in significant fitness costs in the absence of colistin despite providing fitness advantages in the presence of colistin. This suggests that tight control of PET expression is essential, particularly for highly toxic *pet* genes like *eptA* and *mcr-9*. Indeed, *eptA* expression is tightly regulated by two-component systems in *Salmonella* (31). While regulation of *mcr-9* has not been well elucidated, a previous study reported an inverted repeat motif as a putative regulatory element found upstream of 95% of evaluated *mcr-9* genes (21) that may facilitate regulation. Interestingly, we also found that MCR-9 protein levels were significantly lower than MCR-1 and -3 levels when induced with the same L-arabinose concentration. This suggests that PETs are likely post-transcriptionally controlled through mRNA or protein stability, which has not yet been explored in the context of PETs. Exploring mRNA and protein stability of different PETs and their context-dependent fitness advantages in host, low ion, etc. environments will likely provide us with more insight into this group of enzymes.

We also saw that fitness in the presence of colistin was highly variable for strains expressing EptA or MCR-9. Protein levels in heterologous expression systems have been reported to vary across different cells in the same population (32), suggesting that expression of these two PETs likely generates subpopulations that differ in their relative fitness in this study. Our observations suggest that natural populations of *mcr*-carrying bacteria may engage in microbial bet-hedging (33, 34) to help manage the high fitness costs of some *mcr* variants. This hypothesis is supported by observing heteroresistance in *mcr*-positive wild-type strains, including heteroresistance in *mcr-9* expressing strains (30) and reports of heteroresistance as a bet-hedging strategy in bacteria expressing other antibiotic resistance genes (35).

To expand our understanding of PETs, we also used our expression system to characterize two novel, *mcr-*like PETs. Both genes did not confer col^R^ in *E. coli* - either in MIC or killing assays - and appear phenotypically distinct from bonafide PETs by showing moderate cell viability defects. While both PET-B and -C expressing strains showed fitness cost in the absence of colistin, neither showed fitness advantages in the presence of subinhibitory levels of colistin. As PET-B and -C have predominantly been identified in *P. aeruginosa* (4), these proteins may have specific substrate specificity for lipid A produced by *P. aeruginosa* and, hence, do not confer resistance in *E. coli.* This idea is supported by reports of lipid A structural differences between organisms (22) and the fact that we were unable to detect pEtN-modification of lipid A by PET-B. However, we identified PET-C-mediated pEtN-modification of the 4’-phosphate of lipid A, which is the same residue that MCR-1 and -3 modify. The lack of detectable col^R^ in the PET-C expressing strain could be explained by the reduced function of PET-C in *E. coli,* such that the amount of modified lipid A was not appropriate to create measurable phenotypes. As a different study (36) showed that expression of a PET enzyme related to PET-C in *P. aeruginosa* conferred increased col^R^ and decreased cell viability. Thus, our results suggest caution when using heterologous hosts to detect col^R^ conferred by more diverse *pet* variants.

We performed a comparative structural analysis to understand PET-B’s inability to modify E. coli’s lipid A. While the correlation between the structural variability and the ability of PETs to modify lipid A should be cautiously viewed, it is tempting to speculate on the role of the structurally variable hinge regions in enzyme function. For example, it has been shown that the bridging helix and extended loop play a critical role in the function of EptA by acting as a hinge that offers extensive conformational flexibility between the membrane-bound and the catalytic domains (23, 37, 38). Additionally, a study showed that exchanging the linker region connecting the membrane-bound and the catalytic domains from a non-functioning MCR-3 homolog with a linker region of a functional MCR-3 homolog restored its ability to confer col^R^ (39). Thus, it is likely that structural variability in the bridging helix region might relate to PET-B’s inability to modify *E. coli*’s lipid A.

As several papers have suggested a role for MCR in resistance to different AMPs (10–12), we also tested the sensitivity of PET-expressing strains to a set of cell wall stressors and AMPs. Interestingly, we saw increased EDTA sensitivity in EptA and MCR-9 expressing strains, which may suggest that 1-phosphate pEtN modified lipid A may be more susceptible to EDTA-dependent ion depletion. While our preliminary ZOI experiments did not identify any inhibition by LL-37 or lysozyme for any strains, cecropin A, an AMP produced by insects, (17), showed inhibition of all tested strains - with some PET-expressing strains showing marginally smaller ZOIs than the EV, suggesting possible protective capabilities against cationic AMPs other than colistin. Further AMP sensitivity tests using a wider range of AMP concentrations and different testing methods, e.g., competition assays, (12, 40) may thus be valuable.

Importantly, we show for the first time that PETs stereoselectively modify lipid A in *E. coli*. While MCR-1, MCR-3, and PET-C selectively modify the 4’-phosphate of lipid A, EptA and MCR-9 modify the 1-phosphate. For our canonical PETs, which all come from *Enterobacteriaceae* (21, 41, 42), the modified site seems to correlate with phenotypic outputs of col^R^ and fitness costs. Specifically, 4’-phosphate modification is associated with high levels of col^R^ and low toxicity, whereas the 1-phosphate modification is associated with no change in colistin MIC and high toxicity. While previous studies have shown that other PETs modify different LPS structures (i.e., EptB and EptC, which modify phosphate residues of the inner core), this is the first report of stereoselectivity of enzymes that fall into the EptA/MCR clade. These observations suggest a site specificity of colistin action (i.e., targeting of the 4’-phosphate), which could be exploited to develop improved membrane-targeting antibiotics that may use targets other than the 4’-phosphate. Further experiments examining the positional selectivity of additional PETs, including in other bacterial strains, will help our understanding of both the evolution of col^R^ and the potentially different roles PETs play in bacterial physiology.

Overall, our results indicate that PETs are more phenotypically diverse than previously assumed. The six different PETs tested in this study fall into three different groups based on their phenotypes in *E. coli*: (i) PETs that provide high levels of col^R^ but moderate fitness costs [i.e. MCR-1 and -3], (ii) PETs that provide fitness advantages to subinhibitory levels of colistin but high fitness costs in the absence of colistin [i.e. EptA and MCR-9] and (iii) PETs that provide no col^R^ and moderate fitness costs [i.e. PET-C]. Most importantly, our results suggest that the stereoselectivity of pEtN-attachment to lipid A matters for phenotypic outputs of col^R^ and fitness.

## MATERIALS AND METHODS

### Strains, vectors, and growth conditions

*E. coli* Top10 strains were grown at 37°C, 200 rpm in Difco LB Lennox Broth (LB; Becton, Dickinson and Company [BD]; Franklin Lakes, NJ; cat. #240230). For *E. coli* strains carrying pBAD24 or pUC19 vectors, 100 μg/mL of ampicillin (AMP) was added to maintain the vectors.

### Choosing putative, novel *mcr*-like variants

To test our expression system, we furthermore selected two putative, novel *mcr*-like variants (NCBI accession numbers: IPC1619_RS00685 and Q027_RS12935) for cloning and phenotypic characterization in our heterologous host systems. These two genes were selected based on a phylogenetic study of *mcr* and related genes reported by Gaballa et al. (4); this study predicted five putative, novel groups of *mcr* genes. As two of these groups, groups B and C, contained the largest number of sequences (6 and 13, respectively), we selected to use one sequence from each group for these experiments; the other three groups only had two or three predicted members (4). All putative, novel *mcr*-like genes from groups B and C come from *P. aeruginosa* isolates. We selected the putative, novel *mcr*-like gene IPC1619_RS00685 (referred to as *petB*) from group B because it had the most conserved amino acid sequence, i.e., 100% sequence identity with four of the other predicted members of this class. We selected the putative, novel *mcr*-like gene Q027_RS12935 (referred to as *petC*) from group C as this gene is closest to a possible consensus sequence. In comparison to group B, group C genes showed greater sequence diversity, with a total of eleven amino acids differing between putative MCR-like proteins in group C. For ten of these amino acid residues, Q027_RS12935 has the amino acid that is found in most of the isolates. For one amino acid residue (residue 497) Q027_RS12935 encodes a Serine, which is found in five of the thirteen isolates (the other 8 isolates encode a Glycine instead). IPC1619_RS00685 was isolated from a cystic fibrosis patient, while Q027_RS12935 was isolated from a foot wound, suggesting these putative, novel *mcr* genes may have clinical relevance. Lastly, the geographical isolation location for both isolates is in the US which may make it easier to obtain the isolate for future studies if needed.

### Cloning of canonical *mcr* variants, *eptA* and putative, novel *mcr*-like variants

*mcr-3.1* and *mcr-9.1* with 3x FLAG tags were PCR amplified with pBAD24_mcr3_f, pBAD24_mcr9_f and pBAD24_FLAG_r, respectively, from previously created vectors, pET17b-*mcr-3* and pET17b-*mcr-9* (15), with *mcr-3.1* coming from isolate *S. enterica* FSL R9-3269 and *mcr-9* coming from *S. enterica* FSL R9-3274. Primer information can be found in Table S2. *mcr-1.1* (HTI99_RS00160), *eptA* from susceptible isolate *S. enterica* FSL R9-5409 (WGS available under SRX3031601), *petB* (IPC1619_RS00685), and *petC* (Q027_RS12935) with C-terminal 3x FLAG-tag were synthesized in the pFASTBac1 vector with EcoRI and SalI cut sites by ThermoFisher Scientific’s GeneArt Service (ThermoFisher Scientific; Waltham, MA). *Mcr-3.1*, *mcr-9.1*, vector pBAD24 (American Type Culture Collection [ATCC]; Manassas, VA, cat. #87399) and pFASTBac1 vectors with *mcr-1*, *eptA, petB*, and *petC* genes were digested with restriction enzymes EcoRI and SalI (New England Biolabs [NEB]; Ipswich, MA), purified and ligated with T4 DNA ligase (NEB, cat. #M0202L) according to manufacturer’s recommendations. The ligated vectors were transformed into *E. coli* Top10 (ThermoFisher, cat. #C404010) via heat shock, according to the manufacturer’s recommendations. Transformants were selected for by plating on LB + 100 μg/mL Amp and overnight incubation at 37°C. Constructs were confirmed via Sanger sequencing and subsequent analysis in Geneious Prime (Auckland, New Zealand) using the universal pBAD24 forward and reverse sequencing primers (Table S2).

### Cell viability assessment

Cell viability of *pet* expressing strains was assessed by growing *E. coli* strains at 37°C and 200 rpm shaking in LB + 100 g/mL AMP for 12-18 h, followed by 1:200 back dilution in BBL Mueller-Hinton II broth cation-adjusted broth (MHII, BD cat. #212322) + 100 μg/mL AMP and incubation until an OD_600_ of 0.25-0.4 was reached. Canonical *mcr* variants and *eptA* were induced with final L-arabinose (Sigma-Aldrich; St. Louis, MO, cat #A3256-100G) concentrations of 0, 0.01, 0.02, 0.05, 0.1, or 0.2% (w/V) and grown at 37°C and 200 rpm shaking; putative, novel *mcr*-like variants were induced with final L-arabinose concentrations of 0 and 0.05% (w/V). Before induction and after 1, 4, 12, 22 h of induction 100 μL of each strain were removed. Ten-fold serial dilutions were prepared in phosphate-buffered saline (PBS) and 10 μl volumes were spotted onto LB Broth (BD, cat. #240230) agar plates (prepared with 15 g/L Bacto Agar [BD, cat. #214010]) and incubated at 37°C overnight. Colony forming units (CFU/mL) were calculated from colony counts and log_10_-transformed data was plotted. Experiments were performed in biological triplicates. Results were analyzed by a two-way ANOVA and the emmeans package.

### Susceptibility testing

Susceptibility to colistin and EDTA was tested by broth microdilution (BMD) based on the Clinical and Laboratory Standards Institute (CLSI) guidance (43) with the exception being the use of PlateOne polypropylene 96-well plates (USA Scientific; Ocala, FL; cat. #1837-9610) for colistin MIC assays because colistin has a higher binding affinity to polystyrene (44, 45). For EDTA assays, 96-well clear flat bottom polystyrene microplates (Corning; Corning, NY; cat. #3370) were used. Briefly, strains were grown in LB + 100 μg/mL AMP at 37°C and 200 rpm shaking for 12-18 h and subsequently diluted 1:200 in MHII broth + 100 μg/mL AMP, followed by incubation at 37°C and 200 rpm shaking until cultures reached an OD_600_ of 0.25-0.4. Strains were induced by the addition of different concentrations of L-arabinose ranging from 0.01% to 0.2% (w/V), or filter-sterilized dH2O as a negative control and grown at 37°C and 200 rpm shaking for 1 h after arabinose addition. Strains were subsequently inoculated, at a final concentration of 5 x 10^5^ CFU/mL, into MHII broth containing colistin (0.03 – 32 μg/mL, 2-fold dilutions) (Sigma-Aldrich, cat #C4461-1G) and L-arabinose (0, 0.01, 0.02, 0.05, 0.1, 0.2%) and grown statically for 16-20 h at 35°C. Each canonical *mcr* variant, as well as *eptA*, were tested at every L-arabinose concentration in biological triplicates. Putative, novel *mcr*-like variants were tested at 0 and 0.05% L-arabinose in biological triplicates. EDTA assays were performed with all *pet* expressing strains at 0.05% L-arabinose (w/V) and EDTA (Fisher Bioreagents; Pittsburgh, PA, cat #BP120-1) concentrations ranging from 15.625 μg/mL to 16 mg/mL in 2-fold dilutions in MHII broth.

### Western blot analysis of heterologous protein expression

Western blot analysis was performed as previously described to test whether our heterologous PET-FLAG proteins were expressed (15). Briefly, *E. coli* strains were grown at 37°C, 200 rpm shaking in LB + 100 g/mL AMP for 12-18 h, followed by 1:200 backdilution in MHII broth + 100 g/mL AMP and incubation until an OD_600_ of 0.25-0.4 was reached. Cells were induced with 0.05%, 0.2% (w/V) L-arabinose, or dH_2_O as a negative control for 12 h and collected by centrifugation. Cells were lysed in SDS-PAGE lysis buffer by heating (95°C for 10 min) and sonication as detailed by Schumann et al. (15). Lysed cells were resolved on a 4% to 20% Mini-Protean TGX precast protein SDS-PAGE gel (Bio-Rad Laboratories; Hercules, CA, cat. #4568096), followed by protein visualization with the Bio-Rad ChemiDoc MP Imaging system after exposure UV light for 1.5 min. The proteins were transferred to polyvinylidene difluoride (PVDF) membrane and blocked with TTBS (TBS buffer [50 mM Tris-Cl, 150 mM NaCl, pH 7.5] with 0.1% (v/v) Tween 20) containing 5% Blotting-Grade Blocker (Bio-Rad Laboratories, cat. #170-6404) for 30 min. PVDF membranes were incubated with rabbit anti-FLAG primary antibody (Sigma-Aldrich, cat. #F7425, at 1:500 dilution) diluted in TTBS with 0.5% (w/V) skimmed milk powder, at room temperature (RT) overnight with gentle shaking, followed by three 10 min washes with TTBS. PVDF membranes were incubated with goat anti-rabbit-horseradish peroxidase secondary antibody and subsequently developed and visualized as previously detailed (15) . To analyze levels of PET-FLAG proteins semi-quantitively, the area under the curve was estimated for PET-FLAG bands and all protein bands on the SDS page gels via densitometric analysis of band intensity using the Bio-Rad Image Lab 6.1 software as previously described (15). Data was analyzed using a one-way ANOVA with a post-hoc Tukey test.

### Construction of maximum likelihood phylogeny of *mcr* and *mcr*-like genes

Maximum likelihood (ML) phylogeny was inferred from the nucleotide sequences of *E. coli* K-12 substr. MG1655 *eptA*, *eptB,* and *eptC* genes, representatives of *mcr* families, and the newly identified *petB* and *petC* genes. The coding sequences of *eptA*, *eptB,* and *eptC* were extracted from WGS of the *E. coli* K-12 substr. MG1655 (accession number NC_000913.3). Representative of *mcr*-1 to *mcr*-10 families were selected from *mcr* phylogeny (PMID:37360531); briefly, phylogenetic outlier variants in each *mcr* family were excluded (e.g., *mcr*-2.4 and *mcr*-3.17) and the first reported variant in each family was selected. The coding sequences of *mcr*-1.1 (accession number NG_050417.1), *mcr*-2.1 (accession number NG_051171.1), *mcr*-3.1 (accession number NG_055505.1), *mcr*-4.1 (accession number NG_057470.1), *mcr*-5.1 (accession number NG_055658.1), *mcr*-6.1 (accession number NG_055781.1), *mcr*-7.1 (accession number NG_056413.1), *mcr*-8.1 (accession number NG_061399.1), *mcr*-9.1 (accession number MK070339.1), and *mcr*-10.1 (accession number NG_066767.1), *E. coli eptA*, *eptB* and *eptC*, and the newly identified *petB* and *petC* genes were used to construct back-translated nucleotide multiple sequence alignments (NT_btn_-MSA) using MUSCLE (PMID: 15034147) with the default settings in Geneious version 2019.2.3 (Biomatters, Auckland, New Zealand). The resulting NT_btn_-MSAs were used to construct ML phylogenies with 100 bootstrap replicates via RAxML, using the GTRGAMMA substitution model and default settings (RAxML GUI version 2.0.10 and RAxML version 8.2.12) (PMID: 24451623) {Stamatakis, 2014 #107}. The resulting trees were visualized and edited using iTOL version 6.5 (https://itol.embl.de/) (PMID: 17050570).

### Sample Collection for Mass Spectrometry

*E. coli* strains with the pBAD24 vectors were grown at 37°C and 200 rpm shaking in LB + 100 μg/mL AMP for 12-18 h, followed by 1:200 backdilution in MHII broth + 100 μg/mL AMP and incubation until an OD_600_ of 0.25-0.4 was reached. Canonical *mcr* variants and *eptA* were induced with final L-arabinose concentrations of 0, 0.05, or 0.2% (w/V) and grown at 37°C and 200 rpm shaking; putative, novel *mcr*-like variants were induced with final L-arabinose concentrations of 0 or 0.05% (w/V). Cells were collected in 1.5 mL volumes before induction and 1, 4, 12, and 22 h post-induction by centrifugation at 8,000 rpm for 3 min (Eppendorf 5417C centrifuge) and in 3 mL volumes 4 h post-induction by centrifugation at 7,197 x g for 5 min (Sorvall X4RF PRO-MD centrifuge). Putative, novel *mcr*-like variants were also collected prior and after 1-, 2-, 4-, and 12-hours addition of colistin (0.0625 μg/mL). Cells were collected in 1.5 mL volumes (0, 1, 2 hours) and 3 mL volumes (4 and 12 hours), respectively, by centrifugation at 8,000 rpm for 3 mins (Eppendorf 5417C centrifuge). The supernatant was decanted, and pellets were stored at -80°C and shipped to Dr. Robert Ernst on dry-ice.

### Mass Spectrometry

A Bruker microFlex and MALDI (tims TOF) MS were used for FLAT and FLAT^n^ experiments (20, 46). MALDI (tims TOF) MS equipped with a dual ESI/MALDI source with a SmartBeam 3D 10 KHz frequency tripled Nd:YAG laser (355 nm). The system was operated in “qTOF” mode (TIMS deactivated). Ion transfer tuning was used with the following parameters: Funnel 1 RF: 440.0 Vpp, Funnel 2 RF: 490.0 Vpp, Multipole RF 490.0 Vpp, is CID Energy: 0.0 eV, and Deflection Delta: - 60.0 V. Quadrupole has been used with the following values for MS mode: Ion Energy: 4.0 eV and Low Mass 700.00 m/z. Collision cell activation of ions used the following S-4 values for MS mode. Collision Energy: 9.0 eV and Collision RF :3900.0 Vpp. In the MS/MS mode, the precursor ion at m/z 1919.22 was chosen. Isolation width and collision energy were set to 4 m/z and 110 eV, respectively. Focus Pre TOF used the following values for Transfer Time 110.0 µs and Pre Pulse Storage 9.0 µs. Agilent ESI Tune Mix was used to perform calibration of the m/z scale. MALDI parameters in qTOF were optimized to maximize intensity by tuning ion optics, laser intensity, and laser focus. All mass spectra were collected at 104 µm laser diameter with beam scan on using 800 laser shots per spot and 80% laser power, respectively. Both MS and MS/MS data were collected in negative ion mode. A microFlex was used as a comparison to MALDI (tims TOF) MS. In all cases, 10mg/mL of norharman (NRM)1,4 in 1:2 MeOH:CHCl3 (v:v) was used for lipid A detection. NRM solution (1 µL) was deposited on the sample spot. Data Processing: All MALDI (timsTOF) MS and MS/MS data were visualized using mMass (Ver 5.5.0).5 Peak picking was conducted in mMass. Identification of all fragment ions were determined based on Chemdraw Ultra (Ver10.0).

### Killing Assays

*E. coli* Top10 strains with the pBAD24 vectors were grown at 37°C and 200 rpm shaking in LB + 100 μg/mL AMP for 12-18 h, followed by 1:200 backdilution in MHII broth + 100 μg/mL AMP and incubation until an OD_600_ of 0.25-0.4 was reached. All *pet-*expressing strains were induced with a final concentration of 0.05% (w/V) of L-arabinose for 1 h at 37°C and 200 rpm shaking. Cultures OD_600_’s were adjusted to 0.6-0.8, followed by splitting each culture into five, and colistin was added at final concentrations of 0, 1, 2, 4, or 8 μg/mL; these cultures were further grown statically at 35°C for 1 h. Afterwards, ten-fold serial dilutions were prepared in PBS and 10 μl volumes were spot plated onto LB agar plates and incubated at 37°C overnight. Colony forming units (CFU/mL) were calculated from colony counts and the relative log reduction in comparison to the control (grown in 0 μg/mL of colistin) was calculated for each strain and colistin concentration. Experiments were performed in biological triplicates.

### Competition Assays

*E. coli* Top10 (“WT”) was used as a control to calculate the relative fitness of each PET-expressing strain. *E. coli* Top10 strains were competed against *E. coli* Top10 pUC19, which produces blue pigment when grown on media containing 5-bromo-4chloro-3-indoyl β-D-galactopyranoside (X-Gal) (Gold Biotechnology; Olivette, MS, cat # X4281C10). *E. coli* Top10 strains were grown at 37°C and 200 rpm shaking for 12-18 h, followed by 1:200 back dilution in MHII broth + 100 μg/mL AMP for pBAD24 and pUC19 carrying strains until an OD_600_ of 0.25-0.4 was reached. All strains were induced with a final concentration of 0.05% (w/V) of L-arabinose for 1 h at 37°C and 200 rpm shaking. *E. coli* Top10 and *E. coli* Top10 pBAD24 strains were added to fresh MHII media containing 0.05% L-arabinose at a 1:1 ratio with competitor strain *E. coli* Top10 pUC19 at a final concentration of 2.5 x10^5^ CFU/mL. We plated 10-fold serial dilutions of all strains before the competition to determine initial concentrations. Strains were incubated for 12 h at 37°C and 200 rpm, followed by plating 10-fold serial dilutions made in PBS on LB plates supplemented with 20 μg/mL of X-Gal. Plates were incubated overnight at 37°C and white colonies and blue colonies were counted in the countable range (20 – 200 colonies/plate). We used the following formula to calculate fitness:

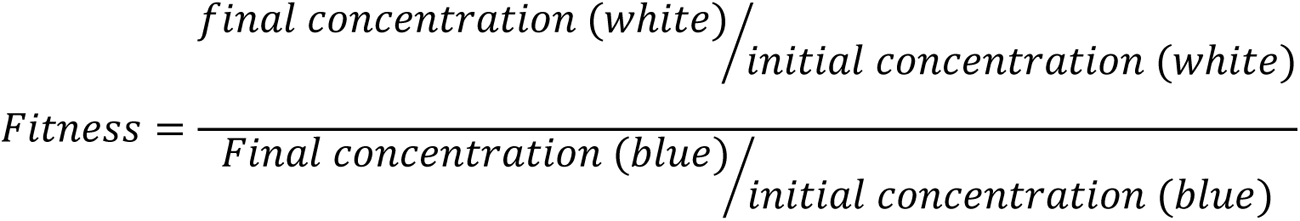

We chose not to adjust our value by log10 or ln to prevent confusion by negative values due to a reduction in cell numbers over the course of the assay for some of our strains. For competitions in the presence of colistin, cells were washed in PBS before the competition assay and colistin was added to the MHII media at concentrations of ¼ of the strain’s MIC (i.e., 0.0625 μg/mL for WT, pBAD24, pBAD24-*eptA*, -*mcr-9*, *-petB*, and *-petC*, 0.5 μg/mL for pBAD24-*mcr-3*, and 1 μg/mL for pBAD24-*mcr-1*). Competition assays were performed in biological quadruplicates and quintuplicates in the absence and presence of colistin, respectively. For the competition assays in the presence of colistin, we added one colony to each count because we could not recover cells for some of the assays. Results were log-transformed, and means were compared by a one-way ANOVA with post-hoc Dunnett’s test or Kruskal-Wallis test with post-hoc Many-to-One Dunn’s test. Variances were compared by calculating the absolute differences between log-transformed values and their group’s median, followed by a one-way ANOVA with a post-hoc Tukey test.

### Zone of Inhibition Assays

For the zone of inhibition assays, MHII plates containing 0.05% L-arabinose were prepared. *E. coli* Top10 pBAD24 strains were grown overnight in LB + 100 μg/mL AMP at 37°C, 200 rpm shaking, diluted 1:20 in 3 mL of MHII Top agar (0.75% agarose) containing 0.05% L-arabinose and spread on MHII plates. Disks made from autoclaved filter paper (Whatman; Maidstone, UK, cat #1001090) were added to plates with sterile forceps and the following compounds were added in 10 μL volumes with final concentrations listed in brackets: colistin (10 μg) [Sigma-Aldrich, cat C4461-1G], nisin (10 μg) [Sigma-Aldrich, cat #N5764-5G], LL-37 (10 μg) [Sigma-Aldrich, cat #94261-1MG], cecropin A (10 μg) [Sigma-Aldrich, cat #C6830-.5MG], lysozyme (10 μg) [Sigma-Aldrich, cat #62971-10G-F}, sodium dodecyl sulfate (1%) [J.T. Baker; Phillipsburg, NJ, cat #4095-02], distilled water (n/a), distilled water pH 2 (n/a). Plates were incubated for 24 h at 35°C and zones of inhibition were measured and recorded. Experiments were performed with biological duplicates.

### Protein Structural Modeling

Structural modeling of PET proteins was done using AlphaFold2 (47) as implemented in the ColabFold v1.5.5 (48) using the default setting and the structure of the *Neisseria meningitidis* phosphoethanolamine transferase EptA structure as a template (38). The structures were viewed and annotated using UCSF ChimeraX V 1.6.1 (49). Structural relationships between different PET proteins were performed using the DALI server (50). Structural similarity matrices and cluster dendrograms are based on the DALI Z-score comparisons calculated from a DALI all-against-all analysis (51). Quality assessment and superimposition of protein structures were done using the Swiss-Model Qualitative Model Energy Analysis (QMEAN) server and Structure Comparison tool (52). Superimposed structures were downloaded, viewed, and edited in UCSF ChimeraX V 1.6.1(49).

### Statistical Analysis

Data were analyzed using R Statistical Software (Version 4.1.2; R Foundation for Statistical Computing, Vienna, Austria).

## DATA AVAILABILITY

All raw data files are available on GitHub (https://github.com/as-2635/PETs_colR_tox) or can be made available on request.

## ACKNOWLEDGEMENTS

This work was funded by a Hatch grant under accession number 1023966 from the USDA National Institute of Food and Agriculture and a grant from the Cornell Center for Pandemic Prevention and Response.

